# Suitability of GRK antibodies for individual detection and quantification of GRK isoforms in western blots

**DOI:** 10.1101/2021.10.26.465910

**Authors:** Mona Reichel, Verena Weitzel, Laura Klement, Carsten Hoffmann, Julia Drube

**Affiliations:** Institut für Molekulare Zellbiologie, CMB – Center for Molecular Biomedicine, Universitätsklinikum Jena, Friedrich-Schiller-Universität Jena, Hans-Knöll Straße 2, D-07745 Jena, Germany

**Keywords:** GRKs, antibody specificity, western blot, protein quantification

## Abstract

G protein-coupled receptors (GPCRs) are regulated by GPCR kinases (GRKs) which phosphorylate intracellular domains of the active receptor. This leads to the recruitment of arrestins resulting in desensitization and internalization of the GPCR. Aside from acting on GPCRs, GRKs regulate a variety of membrane, cytosolic, and nuclear proteins not only via phosphorylation but also by acting as scaffold. This multifunctionality is also reflected by their diverse roles in pathological conditions like cancer, influenza infection, malaria, and metabolic disease.

Reliable tools to study GRKs are the key to specify their role in complex cellular signaling networks. Thus, we examined the specificity of eight commercially available antibodies targeting the four ubiquitously expressed GRK2, GRK3, GRK5, and GRK6 in western blot analysis. We thereby identified one antibody that did not recognize its antigen, as well as antibodies that showed unspecific signals or cross reactivity. Thus, we strongly recommend testing any antibody with exogenously expressed proteins to clearly confirm identity of the obtained western blot results.

Utilizing the most suitable antibodies we established the western blot-based, cost-effective, simple tag-guided analysis of relative protein abundance (STARPA). This method allows comparison of protein levels obtained by immunoblotting with different antibodies. Furthermore, we applied STARPA to determine GRK protein levels in five commonly used cell lines revealing differential isoform expression.

## Introduction

The G protein-coupled receptor kinases (GRKs) were discovered as cytosolic, membrane-associated serine/threonine kinases that phosphorylate ligand-activated G protein-coupled receptors (GPCRs) and thereby allow the binding of arrestins and induce desensitization as well as internalization of the receptor (Lohse and Hoffmann 2014). The human genome encodes seven GRKs (GRK 1-7) that are grouped in three subfamilies: the visual GRK subfamily (GRK1 and 7), the GRK2 subfamily (GRK2 and 3), and the GRK4 subfamily (GRK4, 5, and 6). Four of those GRKs, namely GRK2, 3, 5, and 6 are reported to be ubiquitously expressed (Komolov and Benovic 2018).

Today, GRKs are not only known to phosphorylate GPCRs, but also act on other substrates like receptor tyrosine kinases, cytoplasmic kinases (e.g. src-family kinases), and even nuclear proteins (Martini 2008, Gurevich 2012, Penela 2019). Moreover, GRKs were reported to have phosphorylation-independent scaffolding properties (Raveh 2010, Fernandez 2011). All these activities together explain why GRKs are found to be important in many pathological conditions like cancer, influenza infection, malaria, and metabolic disease (Leoratti 2012, Nogues 2018, Yanguez 2018, Murga 2019).

In order to study these actions of GRKs, it is critical to have reliable tools at hand. Therefore, we have created expression constructs for all four ubiquitously expressed human GRKs (GRK2, 3, 5, 6) in various isoforms (**Table 1**), including versions with point mutations rendering them catalytically inactive (“kinase dead”; GRK2-K220R, GRK3-1-K220R, GRK5-K215R, GRK6-1-K215R). We have utilized these expression plasmids to overexpress the GRKs in HEK293 cells and determined the ability of selected commercially available antibodies to detect the proteins.

**Table 1:** List of eight commercially available antibodies examined in this study targeting the ubiquitously expressed human GRK isoforms. Overview of the tested antibodies against each isoform is provided including the supplier’s information, our review of each antibody as well as the dilution used to determine the antibody specificity. Additionally, the number of amino acids as well as the calculated approx. molecular weight for each human GRK isoform are listed. According to our analysis, our preferred antibody for each GRK isoform is highlighted with bold letters.

|  | number of amino acids | approx. molecular weight [kDa] | ordering number | supplier | source species | antigen as described by supplier | species reactivity as described by supplier | Our findings | antibody dilution used |
| --- | --- | --- | --- | --- | --- | --- | --- | --- | --- |
| GRK2 | 689 | 80 | sc-13143 (C-9) | Santa Cruz Biotechnology | mouse monoclonal | raised against amino acids 468-689 of human GRK2 | human, mouse, rat | good detection of endogenous GRK2 in HEK293, some crossreaction with overexpressed GRK3 | 1 : 500 |
| GRK2 | 689 | 80 | CS #3982 | Cell Signaling Technology | rabbit polyclonal | immunizing animals with a synthetic peptide corresponding to the amino-terminal residues of human GRK2 | human, mouse, rat, hamster, monkey | crossreaction with overexpressed GRK3-1, only weak detection of endogenous GRK2 in HEK293 cells | 1 : 1.000 |
| GRK3-1 | 688 | 80 | sc-365197 (C-11) | Santa Cruz Biotechnology | mouse monoclonal | raised against amino acids 646-688 of human GRK3 | human, mouse, rat | does **not detect** overexpressed human GRK3 isoforms - only background band | 1 : 250 |
| GRK3-2 | 575 | 66 |  |  |  |  |  |  |  |
| GRK3-1 | 688 | 80 | CS #80362 (D8G6V) | Cell Signaling Technology | rabbit monoclonal | immunizing animals with a synthetic peptide corresponding to residues surrounding Lys454 of human GRK3 | human, mouse | slight crossreaction with overexpressed GRK6 isoforms | 1 : 250 |
| GRK3-2 | 575 | 66 |  |  |  |  |  |  |  |
| GRK5 | 590 | 68 | sc-518005 (D-9) | Santa Cruz Biotechnology | mouse monoclonal | raised against amino acids 94-157 of human GRK5 | human, mouse, rat | good detection of endogenous GRK5 in HEK293 | 1 : 250 |
| GRK5 | 590 | 68 | VPA00469 KT | Bio-Rad | rabbit polyclonal | synthetic peptide directed towards the middle region of human GRK5 | human, mouse | strong background band, reported to supplier, was then discontinued | 1 : 1.000 |
| GRK6-1 | 576 | 66 | CS #5878 (D1A4) | Cell Signaling Technology | rabbit monoclonal | synthetic peptide corresponding to residues surrounding Glu89 of human GRK6 | human, mouse, rat | some crossreaction with overexpressed GRK5 | 1 : 1.000 |
| GRK6-2 | 589 | 67 |  |  |  |  |  |  |  |
| GRK6-3 | 560 | 64 |  |  |  |  |  |  |  |
| GRK6-1 | 576 | 66 | PB9709 | Boster Biological Technology | rabbit polyclonal | C-terminus of human GRK6 (amino acids 382-417) | human, rat | Strong background around 55kDa - but can detect human isoform 6-4 | 1 : 1.000 |
| GRK6-2 | 589 | 67 |  |  |  |  |  |  |  |
| GRK6-3 | 560 | 64 |  |  |  |  |  |  |  |
| GRK6-4 | 366 | 42 |  |  |  |  |  |  |  |

Taking a closer look the distribution of the so-called ubiquitously expressed GRKs in specific tissues, it becomes apparent that there are nevertheless striking differences in the expression levels when comparing mRNA data (Matthees 2021).

The relationship between mRNA abundance and actual expressed protein level is not always linear and depends on many different factors like availability of components for biosynthesis or proteasomal degradation (Penela 2003, Liu 2016). In standard laboratory procedures, western blot is a commonly used technique to investigate the actual protein level expressed in cells or tissues. Various companies offer antibodies that are advertised to specifically detect certain proteins. Unfortunately, many of those commercially available antibodies cause unspecific background bands in addition to the intended protein that lead to difficult interpretation, or in the worst case, do not detect their target protein at all (Michel 2009, Bordeaux 2010, Egelhofer 2011, Gilda 2015, Rosell 2020, Yang 2021). Here we investigated the ability of eight different anti-GRK antibodies to detect the targeted GRK isoform and possible cross reactivity against other GRK family members.

The utilization of validated isoform-specific antibodies to detect protein levels in cell lines or primary tissues allows the comparison of protein levels of one specific protein, but the direct comparison of expression levels of different proteins or isoforms using antibodies is not possible. The influence of antibody concentration, specificity, and clonality might have unpredictable influence on the result – e.g. a higher concentration of one antibody would result in more signal, although the protein content stays identical. To overcome this problem, we established the cost-effective simple tag-guided analysis of relative protein abundance (STARPA). This method allows the relative quantification of different protein isoforms using western blotting with validated antibodies.

## Methods

Expression pcDNA3 plasmids for human GRK2 (NP_001610.2), GRK3-1 (NP_005151), GRK5 (NP_005299.1), and GRK6-1 (NP_001004106.1), as well as the kinase dead mutants GRK2-K220R, GRK5-K215R and GRK6-1-K215R were previously described (Drube 2021).

The remaining isoforms GRK3-2 (NP_001349707), GRK6-2 (NP_002073.2), GRK6-3 (NP_001004105.1), and GRK6-4 (NP_001351093.1) were established from cDNA isolated from HEK293 cells. The K220R mutant of GRK3-1 was created by site directed mutagenesis of the wild type plasmid. For the N-terminally HA-tagged GRKs, the encoding sequence for the peptide MYPYDVPDYA was inserted before the start codon of the respective GRK. The identity of all plasmids was confirmed by sequencing.

Pubmed accession numbers for cDNA transcripts and proteins of all species are summarized in **Supplementary Table 1**.

All cell lines were regularly checked for mycoplasma infection using the Lonza MycoAlert mycoplasma detection kit (LT07-318) and were found to be negative.

HEK293 (human, DSMZ Germany, ACC 305), HEK293T (human, DSMZ Germany, ACC 635) NIH-3T3 (mouse, DSMZ Germany, ACC 59), COS-7 (hamster, DSMZ Germany, ACC 60), and Rat-1 (rat, ATCC, CRL-2210) cells were cultured in DMEM (Sigma-Aldrich D6429) supplemented with 10 % fetal calf serum (FCS, Sigma-Aldrich F7524) and 1 % penicillin streptomycin mixture (Sigma-Aldrich P0781). CHO-K1 (monkey, DSMZ Germany, ACC 110) cells were cultured in DMEM-F12 (Gibco, 21041-025) supplemented with 10 % FCS and 1 % penicillin streptomycin mixture. The cell lines HeLa (human, DSMZ Germany, ACC 57), K562 (human, DSMZ Germany ACC 10), and Molm-13 (human, DSMZ Germany ACC 554) were cultured in RPMI1640 (Sigma-Aldrich R8758) supplemented with 10 % heat-inactivated FCS and 1 % penicillin streptomycin mixture. Our HEK293 derivative with knockout of GRK2, 3, 5, and 6 (ΔQ-GRK) is described in (Drube 2021).

For the antibody specificity testing, 7 x10^5^ HEK293 cells were seeded in each well of a 6-well plate and were transfected with 2 μg of respective plasmid DNA using PEI reagent (Sigma-Aldrich, 408727). After 24 hours, cells were washed once with cold phosphate buffered saline (PBS, Sigma-Aldrich, P4417), and lysed with RIPA lysis buffer (1 % NP-40, 1 mM EDTA, 50 mM Tris-HCl pH 7.4, 150 mM NaCl, 0.25 % sodium-deoxycholate) including protease (Roche 04693132001) and phosphatase inhibitors (Roche 04906845001) diluted in RIPA buffer according to the manufacturer’s recommendations. For the experiments without phosphatase inhibitors, the RIPA buffer was prepared without EDTA and only supplemented with protease inhibitors diluted in this EDTA-free buffer. To the cleared lysates, 20 μl of 6 X sample buffer (375□mM Tris-HCl pH 6.8, 12□% SDS, 30□% glycerol, 500 mM DTT) were added per 100 μl lysate. Samples were boiled for 5 minutes at 95 °C and equal amounts of the lysates were loaded onto polyacrylamide gels and blotted onto nitrocellulose membranes. The membranes were blocked with 5 % dry milk in 1 X TBST (10 X TBST: 200 mM Tris-HCl pH 7.6, 1.37 M NaCl, 10 ml Tween 20), and incubated in the diluted primary antibodies (all diluted in 5 % bovine serum albumin in 1 X TBST) over night at 4 °C with gentle shaking. The utilized GRK antibodies are listed in **Table 1**, actin antibody was purchased from Sigma-Aldrich (A5441, dilution 1:5 000), vinculin antibody was obtained from Biozol (BZL03106, dilution 1:1 000). After washing with 1 X TBST, the membranes were incubated in 5 % dry milk in 1 X TBST of the respective secondary HRP-coupled antibody (sera care, anti-rabbit #5220-0336, anti-mouse #5220-0341, both 1:10 000) and analyzed using a Fuji LAS4000.

For the quantitative analysis, 5 x 10^6^ ΔQ-GRK HEK293 cells were seeded per 10 cm dish and transfected with 20 μg of the respective HA-tagged GRK expression plasmid using PEI reagent. Cells were lysed with 1 ml of RIPA buffer (containing phosphatase and protease inhibitors) and processed as described above. The obtained lysates were supplemented with sample buffer as described above, and further diluted with 1 X sample buffer where indicated. Multiple western blots were performed as described above using an anti-HA antibody (Cell signaling technology #3724, dilution 1:1 000). Quantification was done using the Fujifilm Multi Gauge software V3.0.

For analysis of the endogenous GRK expression, cells were cultivated in full medium, washed with cold PBS, and lysed in RIPA containing protease and phosphatase inhibitors. The total protein concentration of the cleared lysates was determined using a BCA Protein-Assay (Thermo Scientific 23225) and 20 μg of protein was loaded per lane and further processed as described above.

## Results

Lysates with overexpression of GRK2, GRK3-1, GRK3-2, GRK5, GRK6-1, GRK6-2, GRK6-3, GRK6-4 were prepared and analyzed using eight different commercially available GRK antibodies as listed in **Table 1**.

For GRK2, two different antibodies were tested (**Figure 1a, b**). The Santa Cruz Biotechnology antibody raised against GRK2 (**Figure 1a**) detects overexpressed GRK2 and endogenous levels of GRK2 with some detection of GRK3-1. The Cell Signaling Technology antibody #3982 (**Figure 1b**) is also able to detect overexpressed GRK2, but the signal for endogenously expressed GRK2 is weak. This antibody detects overexpressed GRK3-1 with a stronger signal in comparison to the Santa Cruz antibody. The differences in the cross reactivity of these two different GRK2 antibodies with the GRK3-1 protein are very surprising, as the alignment of the GRK2 epitope regions compared to the GRK3-1 sequence shows identical amino acids in 78 % (Santa Cruz antibody, **Supplementary Figure 1a**) or 77 % (Cell Signaling Technology antibody, **Supplementary Figure 1b**). While the similarity between these two GRK isoforms can explain the occurrence of cross reactivity, the relative difference in the extent of cross reactivity cannot be explained by this alignment.

**Figure 1:**
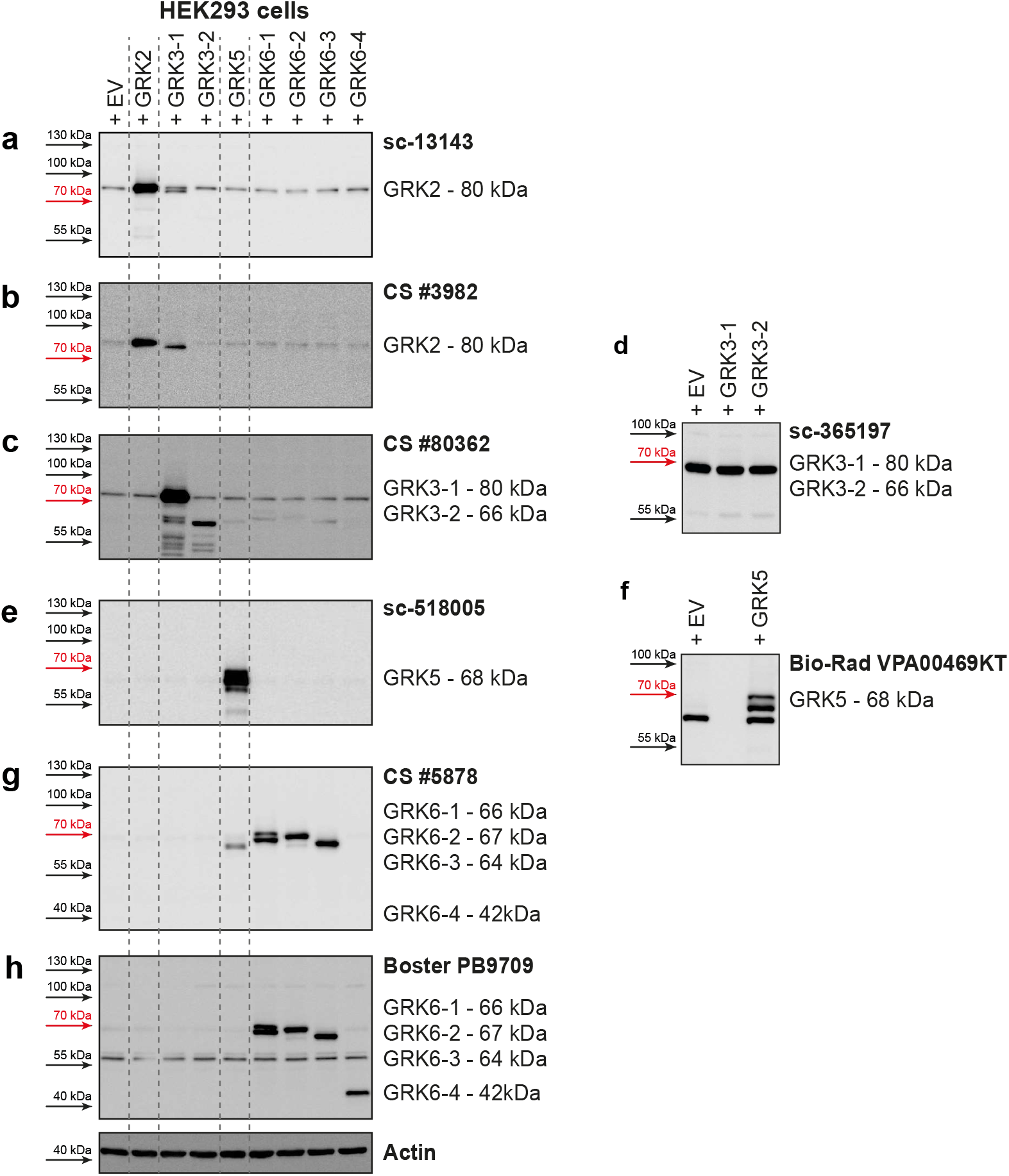
GRK isoform specificity of several commercially available GRK antibodies. HEK293 cells were transiently transfected with pcDNA3 plasmids encoding either GRK2, GRK3-1, GRK3-2, GRK5, GRK6-1, GRK6-2, GRK6-3, or GRK6-4 as indicated. GRK isoform specificity of indicated GRK antibodies was investigated in western blot analysis. Representative blots are shown. Two different antibodies raised against GRK2 (**a, b**), GRK3 (**c, d**), GRK5 (**e, f**), and (**g, h**) were used for the analysis. Approximate molecular weight of each antigen is stated. Detailed information on each antibody examined is listed in **Table 1**. Actin served as loading control.

The overexpression of GRK3-1 and GRK3-2 was detected by the Cell Signaling Technology antibody #80362 (**Figure 1c**). This antibody also mildly detects overexpressed GRK5, GRK6-1, 6-2, and 6-3 isoforms although only 34-35 % of the amino acids match (**Supplementary Figure 1c**). The Santa Cruz Biotechnology antibody sc-365197 was not able to detect the overexpressed GRK3 isoforms, but it gave a strong background band in all HEK293 lysates slightly below the 70 kDa marker (**Figure 1d**).

Overexpressed GRK5 protein is strongly detected by Santa Cruz Biotechnology sc-518005 antibody with no visible cross reactivity to overexpressed GRK6 isoforms (**Figure 1e**). A second tested antibody from Bio-Rad (VPA00469KT) was also able to detect overexpressed GRK5 but additionally displayed a strong background band slightly below the specific protein band (**Figure 1f**). Our findings were reported to the supplier and the antibody was discontinued.

We created expression plasmids for four different GRK6 isoforms. GRK6-1, GRK6-2, and GRK6-3 only differ in the 30 amino acids on the C-terminus, and GRK6-4 is a 210 amino acid N-terminally truncated version of GRK6-1. We tested two antibodies for this GRK. The antibody from Cell Signaling Technologies (#5878) raised against the N-Terminus of GRK6-1, which is identical in GRK6-2 and 6-3, was able to detect overexpressed proteins of these isoforms but was unable to detect GRK6-4 (**Figure 1g**) as this isoform does not include the epitope of this antibody. We observed a slight cross reactivity with overexpressed GRK5 which has 59 % similarity in the amino acid sequence surrounding glutamate 89 of GRK6, which was used as antigen for antibody generation according to the supplier (**Supplementary Figure 1d**). The second tested antibody from Boster Biological Technology (PB9709) was raised against the amino acids 382-417 of GRK6. This region is also present in GRK6-4. As expected, this antibody detected also GRK6-4 in addition to the isoforms GRK6-1, 6-2, and 6-3. However, it gave a strong unspecific band around 55 kDa (**Figure 1h**).

The western blots of overexpressed GRK5 and GRK6-1 (**Figure 1e-h**) revealed double bands of the proteins. Notably, GRKs are known to be (auto)phosphorylated (Palczewski 1997, Pronin and Benovic 1997, Penela 2019). To clarify whether the identified size shift was caused by phosphorylation of the GRKs, we transfected HEK293 cells with either GRK2, 3, 5, and 6 isoforms or with the catalytically inactive variants GRK2-K220R, GRK3-1-K220R, GRK5-K215R, and GRK6-1-K215R. These cells were then lysed in presence of phosphatase inhibitors (+) or in absence of EDTA and phosphatase inhibitors (-) to allow endogenous phosphatases to dephosphorylate the proteins while preparing the lysates. In samples overexpressing GRK2 (**Supplementary Figure 2a**) or GRK3 (**Supplementary Figure 2b**), we could not detect an influence of the endogenous phosphatases nor the expression of the kinase dead mutants on the western blot pattern. The western blot analysis of GRK5 overexpressing cells revealed, that the upper band visible in the lysates of wild type GRK5 treated with phosphatase inhibitors is due to phosphorylation of the kinase since the band is strongly diminished in the lysates with the active phosphatases (**Supplementary Figure 2c**). The kinase dead mutant does not show any change with or without active phosphatases. Similar findings are obtained in case of GRK6-1 with both tested antibodies (**Supplementary Figure 2d, e**). These findings indicate a strong phosphorylation of GRK5 and 6 in our experimental setup, leading to a size shift.

The antibodies are described to detect the respective GRK not only in humans, but also in other species like mouse, hamster, rat, and monkey (**Table 1**). Therefore, we tested the ability of the GRK antibodies to detect endogenously expressed GRKs in HEK293 (human), NIH-3T3 (mouse), COS-7 (hamster), Rat-1 (rat), and CHO-K1 (monkey) cell lines. All these species express different GRK isoforms homologous to the human GRKs (**Supplementary Table 1**). Both GRK2 antibodies are able to clearly show a signal corresponding to the size of human GRK2 in all analyzed cell lines (**Figure 2a, b**). The amino acid sequence alignment in the respective regions of antibody recognition shows 96 % to 100 % identity to the human GRK2 sequence (**Supplementary Figure 2a, b**).

**Figure 2:**
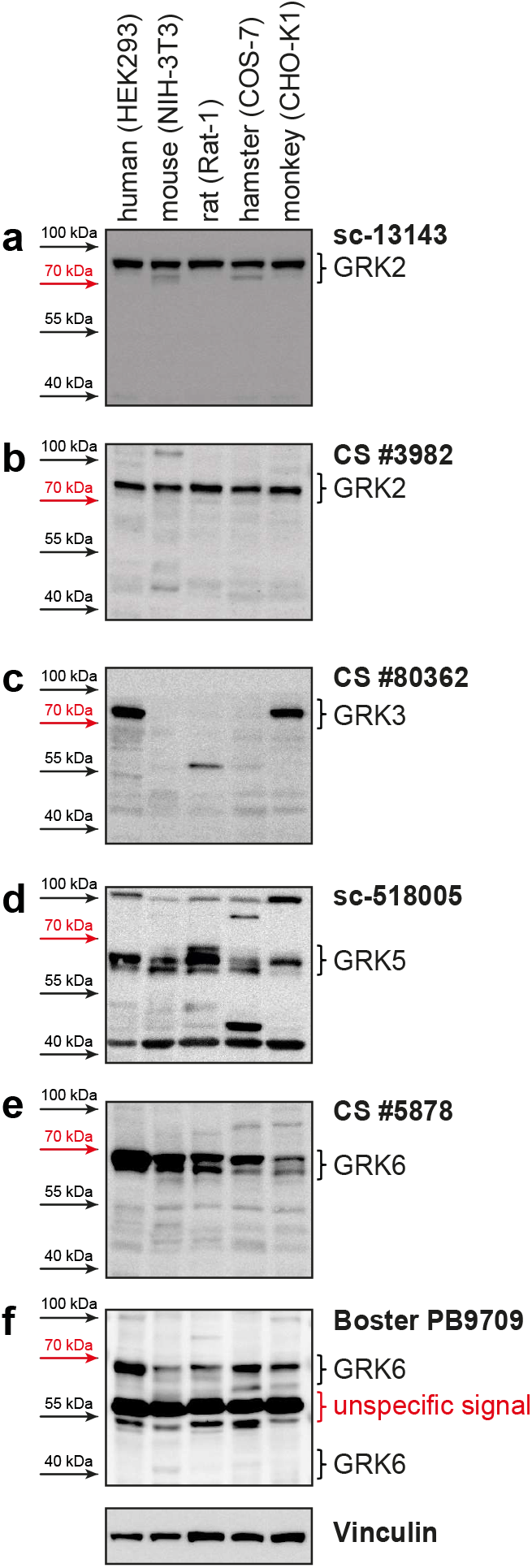
**a-f** Western blot analysis of HEK293 (human), NIH-3T3 (mouse), COS-7 (hamster), Rat-1 (rat), and CHO-K1 (monkey) cell lysates using the denoted anti-GRK antibodies. Vinculin served as loading control.

The GRK3 antibody does only detect the endogenously expressed GRK3 in the human and monkey cell lines (**Figure 2c**). These two species have the identical amino acid sequence in the antibody detection region. The amino acid sequence of mouse, rat, and hamster GRK3 however are more variable with only 89 %, 88 %, and 90 % identity, respectively (**Supplementary Figure 4**). These differences might explain the absence of a specific GRK3 band. However, we cannot exclude that GRK3 is absent in these analyzed cell lines under our experimental conditions.

Utilization of the GRK5 antibody leads to detectable signals in all tested cell lines (**Figure 2d**) in the expected protein size. The similarity to human GRK5 is 87 – 95% (**Supplementary Figure 5**).

A signal corresponding to the size of human GRK6 can be detected in all analyzed cell lines, with both tested antibodies (**Figure 2e, f**). Notably, there is no clearly visible band reflecting the GRK6-4 isoform (**Figure 2f**). The GRK6 sequences of the different species are very similar to huGRK6-1 – ranging from 92 – 96 % similarity for the epitope region of the Cell Signaling Technology antibody (**Supplementary Figure 6a**) and 88 – 100 % similarity for the Boster Biological Technology antibody (**Supplementary Figure 6b**).

Next, using our quadruple GRK knockout HEK293 cells (ΔQ-GRK, lacking endogenous expression of GRK2, 3, 5, and 6 (Drube 2021)), we established the cost-effective simple tag-guided analysis of relative protein abundance (STARPA). This method allows the relative quantification of GRK protein amount in western blot samples with unknown GRK isoform expression levels. **Figure 3a** and **3b** schematically depict the principal STARPA concept: after transfection of the HA-GRK isoforms, cell lysates are analyzed for their HA-antibody signal and the optimal dilution factor to result in equal signals for all HA-GRK isoforms is experimentally determined by sample dilution (**Figure 3a**). These standardized samples are then loaded onto the same gel as reference in addition to the lysates with unknown GRK expression. Utilizing the GRK-specific antibodies, the normalized (arbitrary units (AU) of unknown sample / AU of standard) signal will reflect the relative amount of this specific isoform (**Figure 3b**).

**Figure 3:**
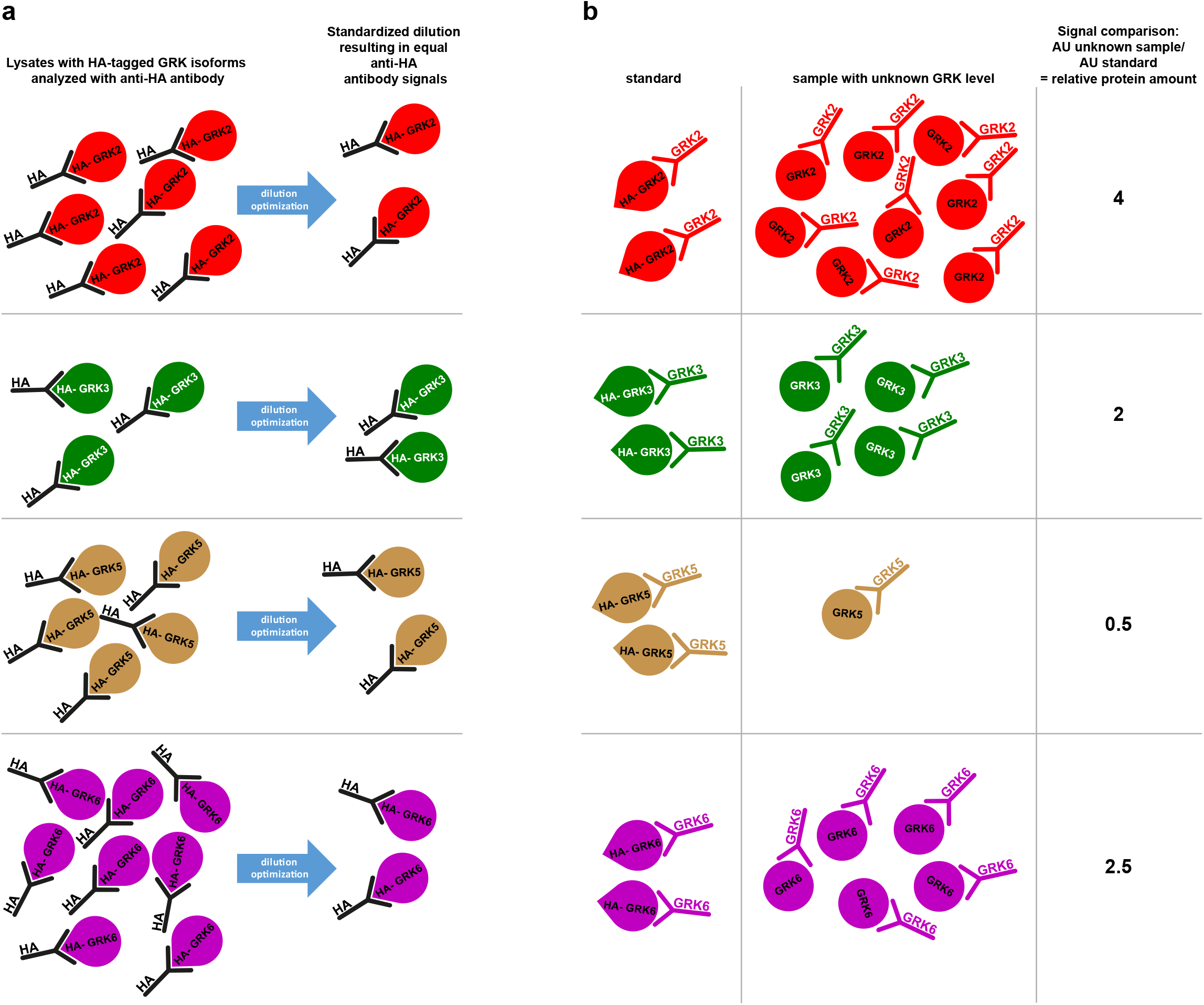
Simple tag-guided analysis of relative protein abundance (STARPA). Schematic depiction of STARPA implementation (a) and functionality (b). The creation of a protein standard for each GRK isoform containing equal amounts of HA-tagged GRK (a) enables the analysis of relative GRK levels via western blot in any sample containing untagged GRK using GRK isoform-specific antibodies (b). AU, arbitrary units

First, we prepared lysates of overexpressed HA-tagged GRK isoforms in ΔQ-GRK cells, as these are devoid of endogenous GRK expression. Therefore, any western blot signal of GRKs would be caused by the overexpressed tagged GRK versions and no purification steps are necessary. We created multiple dilution series ranging from 1:10 to 1:100, and they were analyzed by western blotting using an anti-HA antibody (**Figure 4a**). Quantification of several independently mixed dilutions and multiple analyses of them allowed the calculation of optimal standard dilutions (**Figure 4b**). In our setup, we chose the HA-GRK2 1:20 dilution as reference and calculated all obtained signals for each blot relative to that result in order to minimize differences in developing times. Subsequently, we identified the dilutions of the other GRK isoforms that displayed the smallest differences in HA signals compared to our GRK2 reference dilution, namely GRK3 1:10, GRK5 1:20, and GRK6 1:40. Next, new dilution series for each GRK were prepared for fine-tuned assessment of optimal results (GRK3 1:8, 1:10, 1:12; GRK5 1:18, 1:20, 1:22; GRK6 1:30, 1:35, 1:40, 1:45) and analyzed again by immunoblotting (data not shown). The best matching dilutions were then extensively validated (**Figure 4c, d**).

**Figure 4:**
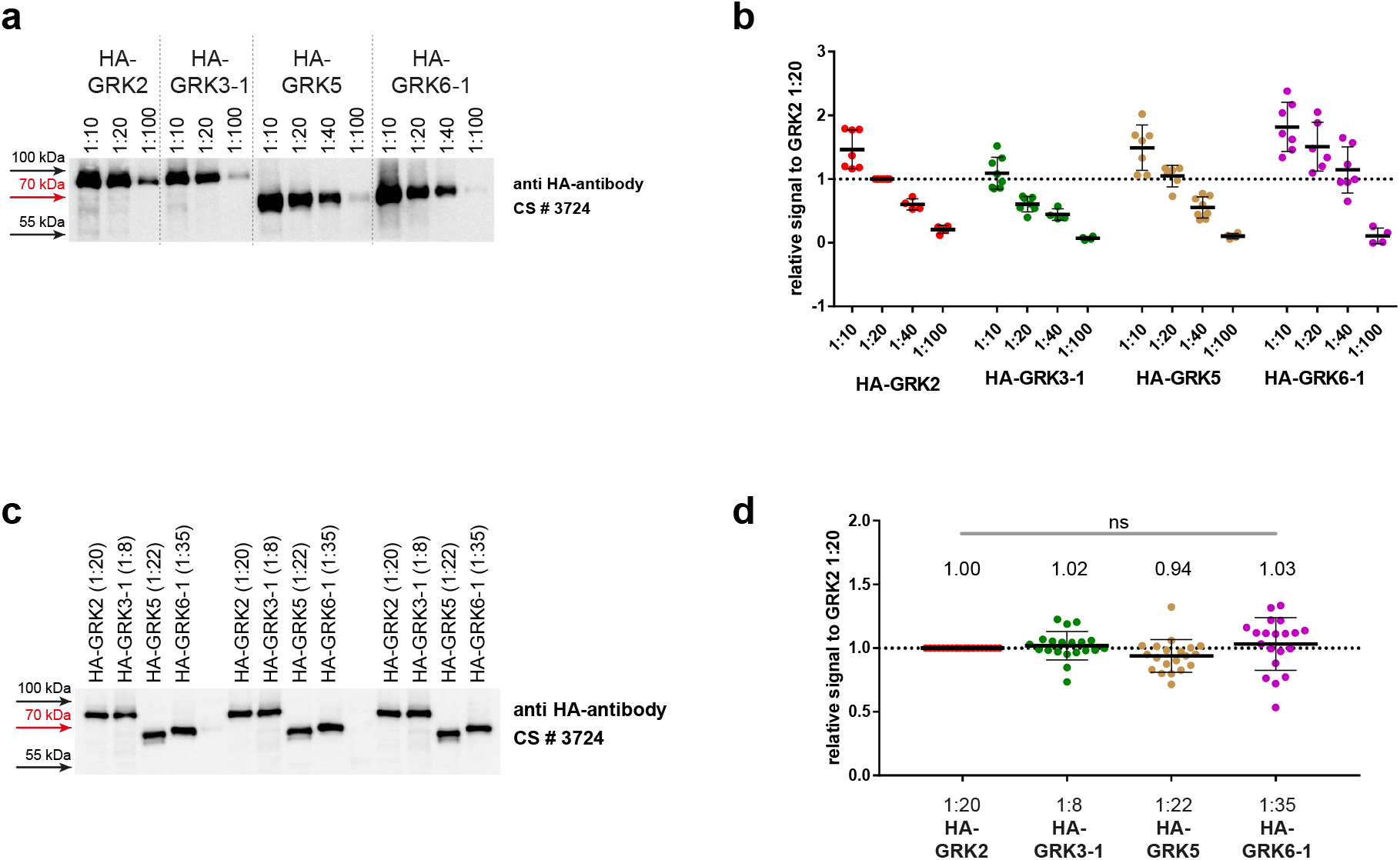
Implementation of protein standards for STARPA. **a** To create the HA-tagged GRK protein standards for STARPA, ΔQ-GRK cells (HEK293 cells with knockout of GRK2, 3, 5, and 6, described in Drube 2021) were transiently transfected with HA-tagged GRK2, 3-1, 5 or 6-1. Serial dilutions of lysates were analyzed for their GRK levels using anti-HA-antibody. A representative blot is shown. **b** Blots were quantified and the relative signal compared to the respective signal in the 1:20 dilution of the HA-GRK2 are shown as dot plots with corresponding means +/- standard deviation (SD), indicated by bars. **c** Accordingly to the results shown in (**b**), new dilutions from the same lysates were created with the aim to carry equal protein concentrations of the corresponding HA-GRK isoform. Their HA-GRK level was determined via western blot analysis using anti-HA-antibody. A representative blot showing 3 of 20 independent datasets is shown. **d** Blots were quantified and the relative signal compared to the respective signal in the 1:20 dilution of the HA-GRK2 are shown as dot plots with corresponding means +/- standard deviation (SD), indicated by bars. Additionally, means are stated above the dot plots. No statistical difference (ns) was found using one-way ANOVA und Tukey’s test.

These validated standards were then loaded onto western blot gels in parallel with 20 μg of total protein from HEK293, HEK293T, HeLa, K562, and Molm-13 lysates with unknown GRK content (**Figure 5a**). The samples were analyzed with GRK specific antibodies as indicated and additionally with anti HA-antibody.

**Figure 5:**
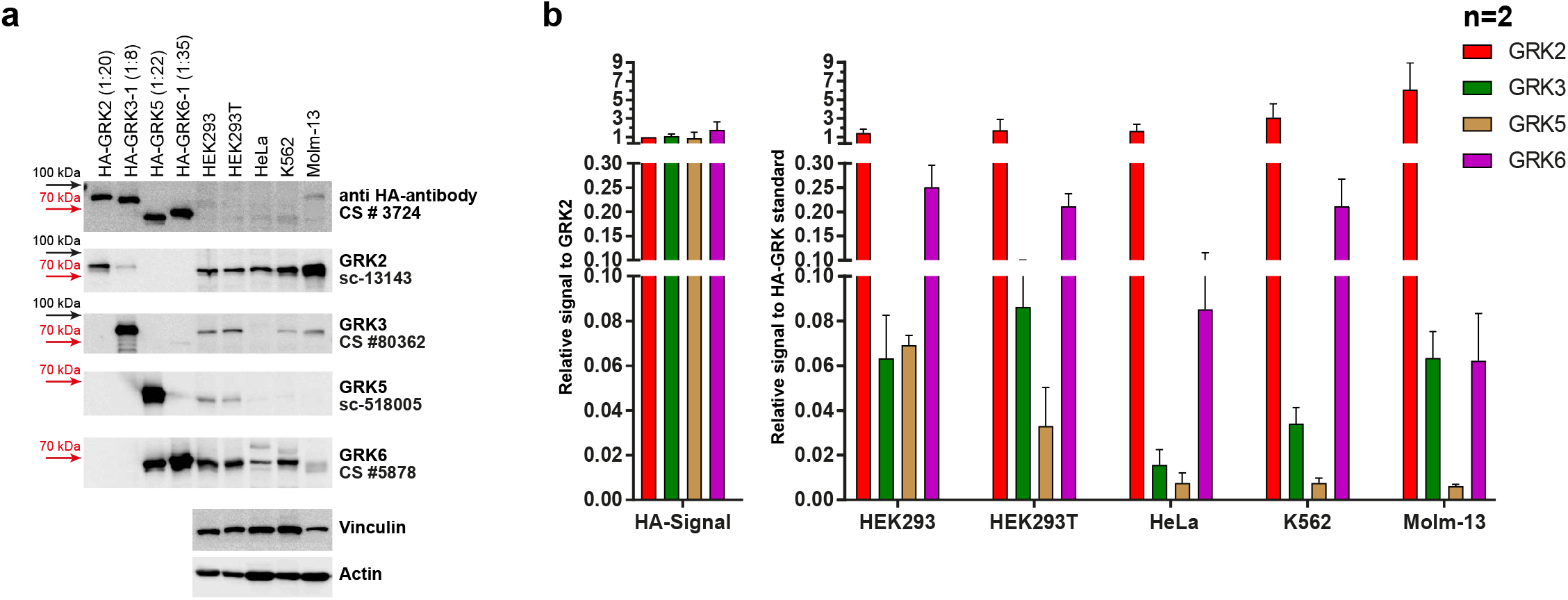
STARPA reveals differential GRK expression in commonly used cell lines. **a** STARPA was used to determine relative GRK expression in HEK293, HEK293T, HeLa, K562, and Molm-13 cells. Lysates of HEK293, HEK293T, HeLa, K562, and Molm-13 cells together with the STARPA standards were loaded onto western blot gels. Western blot analysis using anti-HA-antibody confirmed the GRK protein abundance in the STARPA standards. Immunoblotting using anti-GRK-antibodies for each isoform allows the determination of endogenously expressed GRK levels in the cell lines. Shown are representative images of 2 independent blots from identical lysates. **b** Blots were quantified according to each GRK isoform-specific antibody and the relative signal compared to the respective signal in the standard is shown as mean + SD of two blots from identical lysates.

After quantification of the immunoblots, the cell line signals obtained from incubation with GRK specific antibodies were divided by the signal with the GRK specific antibody of the respective standard, resulting in the relative GRK expression pattern of the analyzed cell lines (**Figure 5b**). GRK2 was the most abundant isoform in all five tested cell lines. In HEK293, HEK293T, HeLa, and K562 cells, GRK6 was the second highest expressed isoform although reaching only 5 – 17 % of their respective GRK2 expression level. Molm-13 cells have a very low expression of GRK6 with only 1 % of the GRK2 expression level. GRK3 and GRK5 were much lower expressed than GRK2 and GRK6 in all cell lines except for Molm-13, in which GRK3 and GRK6 were similarly expressed. The lowest expression was detected for GRK5 in HeLa, K562, and Molm-13 with less than 0.5 % of the respective GRK2 expression. Interestingly, the related cell lines HEK293 and HEK293T show a similar expression level of GRK2, 3, and 6, but the expression of GRK5 is approximately 2-fold higher in HEK293 than in HEK293T.

Taken together, the employment of STARPA demonstrates differential GRK isoform expression in these commonly utilized cell lines.

## Discussion

Most of the tested antibodies (GRK2: sc-13143, CS #3982; GRK3: CS #80362; GRK5: sc-518005, VPA00469KT; GRK6: CS #5878, PB9709) are able to detect the targeted protein, but some also strongly label background bands with similar protein size leading to difficult interpretation of the expression levels especially at endogenous levels (GRK5: VPA00469KT, GRK6: PB9709). One antibody (sc-365197, GRK3) did not detect its target GRK at all, but detected a very strong background band only. In our test setup, we also observed some cross reactivity with the other overexpressed GRK isoforms. We could detect overexpressed GRK3 while using GRK2 antibodies and GRK5 when using GRK6 antibody. This observed cross reactivity should be taken into consideration, as the influence on the obtained result depends on the expression ratio of GRK3 to GRK2 and GRK5 to GRK6. This cross reactivity is also the reason, why the four standards for GRK quantification could not be mixed into one standard containing all GRK isoforms.

As GRKs are conserved among mammals (Premont 1999), the possibility that a given antibody would detect the same GRK in other species is obvious. In order to get an impression, whether the analyzed antibodies would also be able to detect the proteins of mouse, rat, hamster, or monkey, we analyzed cell lines of these species. For GRK2, GRK5, and GRK6, we detected bands in the size comparable to the huGRKs, but in case of GRK3, we could not detect a signal in mouse, rat, or hamster cells, although the supplier states the recognition of mouse GRK3. The absence of detectable protein in the tested cell lines could be either due to the lack of antibody affinity to the species typical GRK, or simply due to the absence of that GRK in the tested cell line. To finally validate the recognition of GRKs from different species, it would be mandatory to test the antibodies with overexpressed isoforms of the species of interest, and/or in protein lysates of knock out (KO) animals or cell lines.

In this study we compared closely related proteins but the proposed STARPA method allows the relative comparison of different expression levels of any protein that can be expressed with the same protein tag. By introduction of the standards, the variability of antibody concentration and affinity can be normalized and thereby allow direct comparison. We used the ΔQ-GRK cell line without detectable GRK2, 3, 4, and 6 expression (Drube 2021) for expression of HA-GRK isoforms. In that case, no purification of the HA-tagged GRKs was needed, as there is no endogenous GRK present. If no KO cells are available to overexpress the desired protein, an immunoprecipitation of the tagged proteins should be carried out before the creation of the dilution series to find optimal diluted samples.

We used our established quantification method to compare the GRK2, 3, 5, and 6 expression in different cell lines. The here reported differences and expression patterns reflect a snapshot of the GRK levels in our growth conditions. GRK levels can change during progression of cell cycle (Penela 2010) or the expression pattern might be influenced by the level of nutrients, cell density, plasticware, cell passage, and other factors (Geraghty 2014). Taken all these factors into account, the expression level of a given cell line might be varying between different laboratories and must be individually checked. Our analysis demonstrates the ability of STARPA to reveal the differential protein expression of the GRK isoforms in cells derived from various tissues.

Taken together we strongly recommend testing any antibody with exogenously expressed proteins to clearly confirm identity of the obtained western blot results. Furthermore, we propose STARPA as a cost-effective method to compare relative levels of different proteins which can be conducted using standard laboratory equipment.

## Author contribution

M.R. and J.D. developed the concept, conducted experiments, and wrote the manuscript, V.W. and L.K. conducted experiments, C.H. wrote the manuscript

---

**Supplementary Figure 1:**
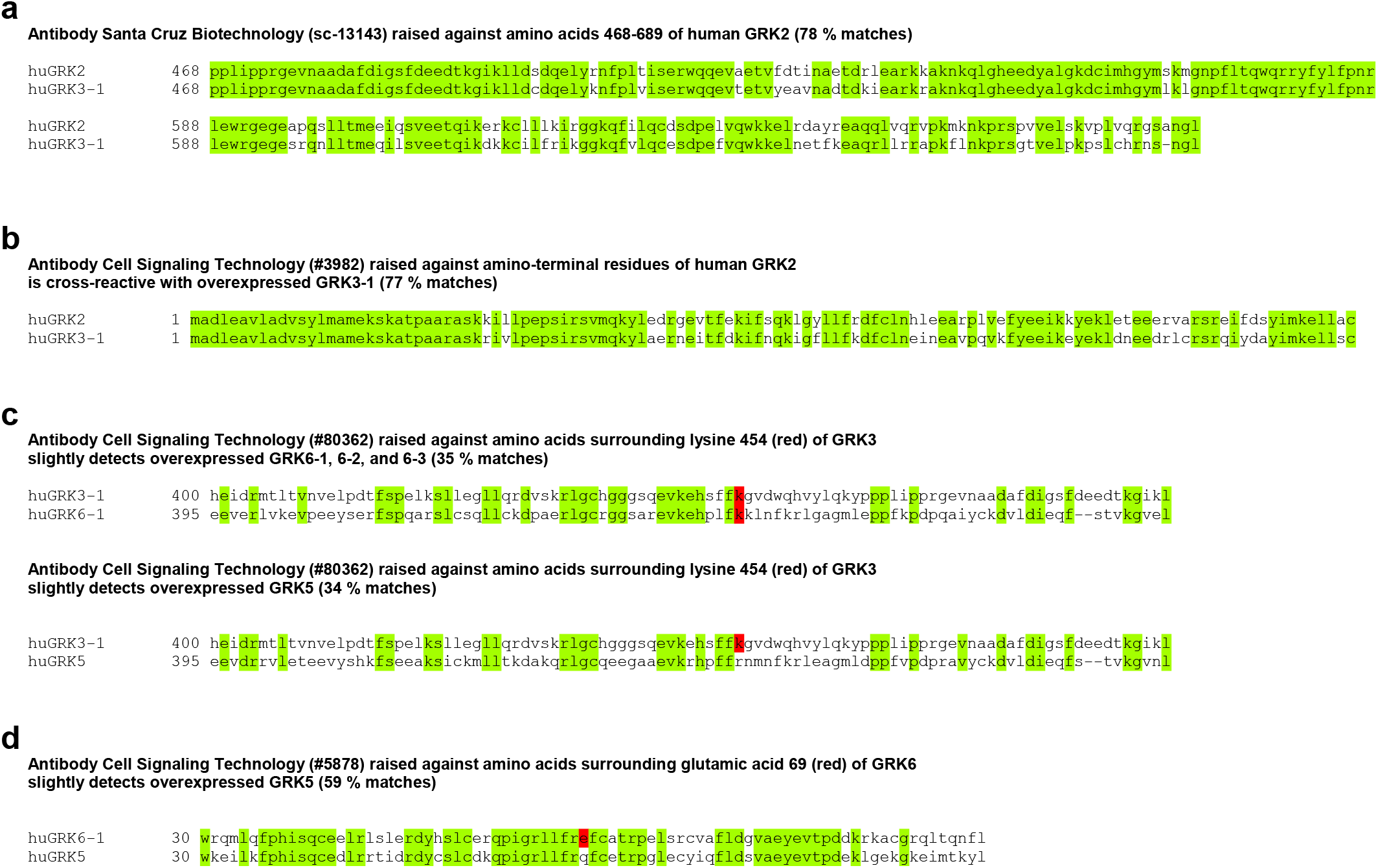
Alignment of protein sequences used to create designated antibodies and the corresponding sequence of the indicated GRK isoform which are also detected by the antibody. Amino acids identical in both sequences are highlighted in green. Total identity is denoted in per cent. a Alignment of human GRK2 and human GRK3-1 protein sequences from amino acid 468 to 689 which are both recognized by antibody sc-13143 (Santa Cruz Biotechnology). b Alignment of human GRK2 and human GRK3-1 protein sequences from the first amino acid to amino acid 120 which are both recognized by antibody CS #3982 (Cell Signaling Technology). c Alignment of human GRK3-1 with human GRK6-1 and GRK5 protein sequences from amino acid 400 to 499 and 395 to 494, respectively. Antibody CS #80362 (Cell Signaling Technology) was raised against GRK3 but also detects GRK6-1, −2, −3 and GRK5. d Alignment of human GRK6-1 and human GRK5 protein sequences from amino acid 30 to 499 which are both recognized by antibody CS #5878 (Cell Signaling Technology).

**Supplementary Figure 2:**
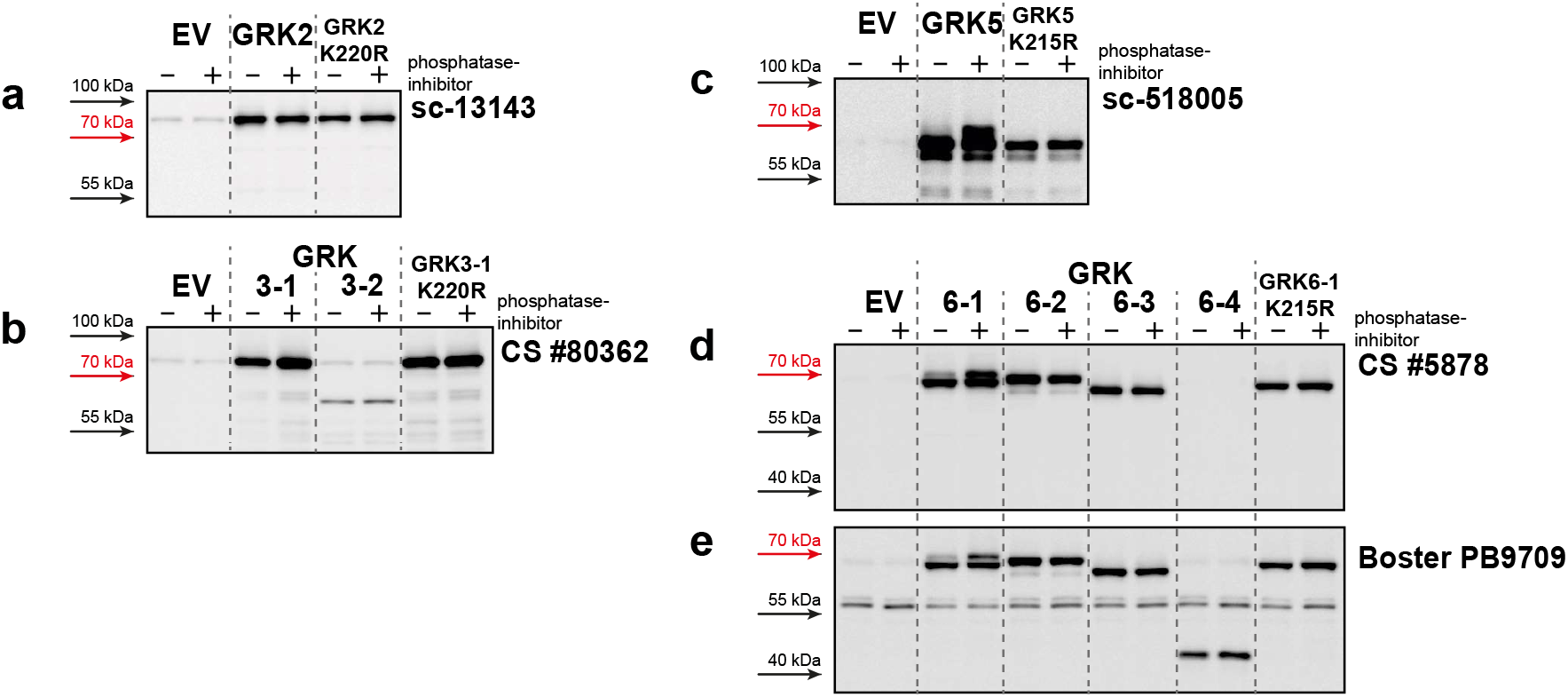
HEK293 cells were transfected with either empty vector (EV), GRK2 (a), 3 (b), 5 (c), and 6 isoforms (d) or their corresponding kinase-dead variants GRK2-K220R (a), GRK3-1-K220R (b), GRK5-K215R (c), and GRK6-1-K215R (d). Lysates were prepared in presence (+) or absence (-) of phosphatase inhibitors and EDTA. Western blot analysis of the lysates using the indicated antibodies was performed.

**Supplementary Figure 3:**
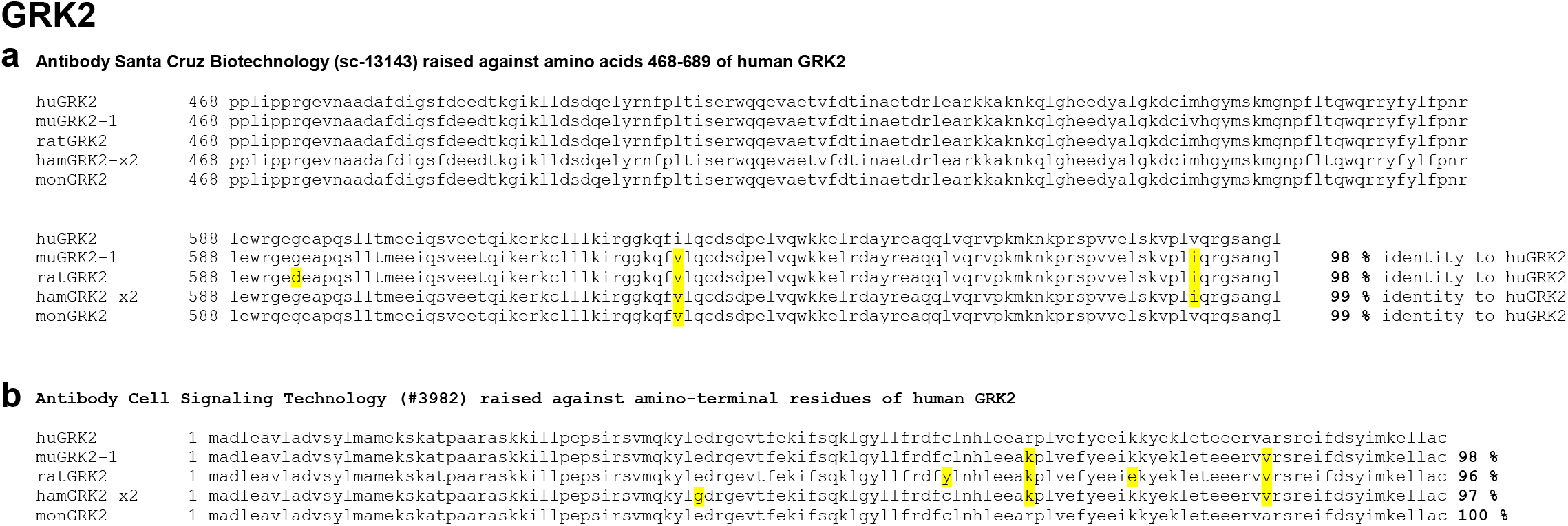
Alignment of human, mouse, rat, hamster, and monkey GRK2 protein sequences from amino acid 468 to 689 (**a**) and 1 to 120 (**b**). Differing amino acids are highlighted in yellow. Total identity is denoted in per cent.

**Supplementary Figure 4:**
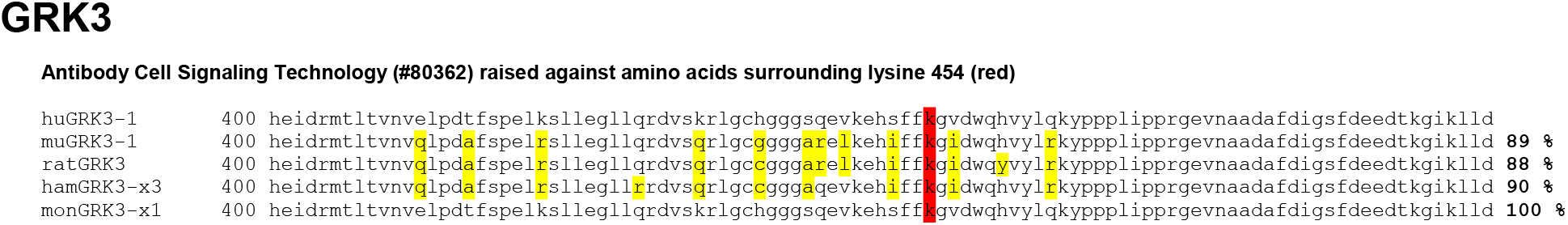
Alignment of human, mouse, rat, hamster, and monkey GRK3 protein sequences from amino acid 400 to 100. Differing amino acids are highlighted in yellow. Total identity is denoted in per cent.

**Supplementary Figure 5:**
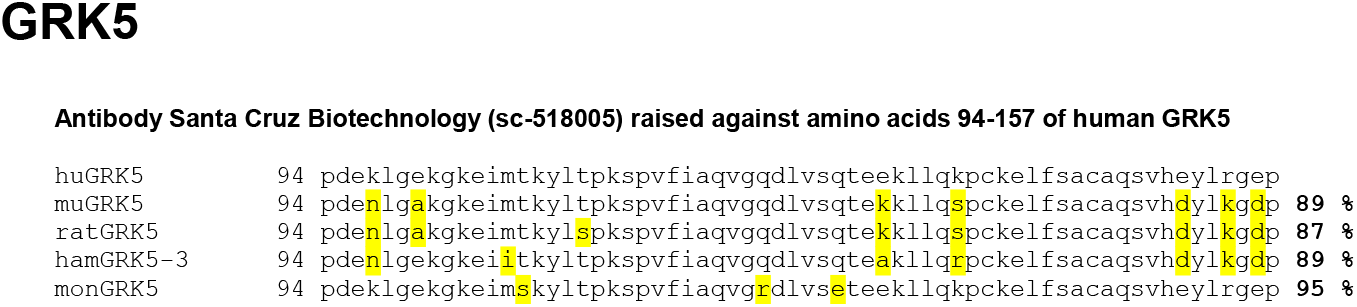
Alignment of human, mouse, rat, hamster, and monkey GRK5 protein sequences from amino acid 94 to 157. Differing amino acids are highlighted in yellow. Total identity is denoted in per cent.

**Supplementary Figure 6:**
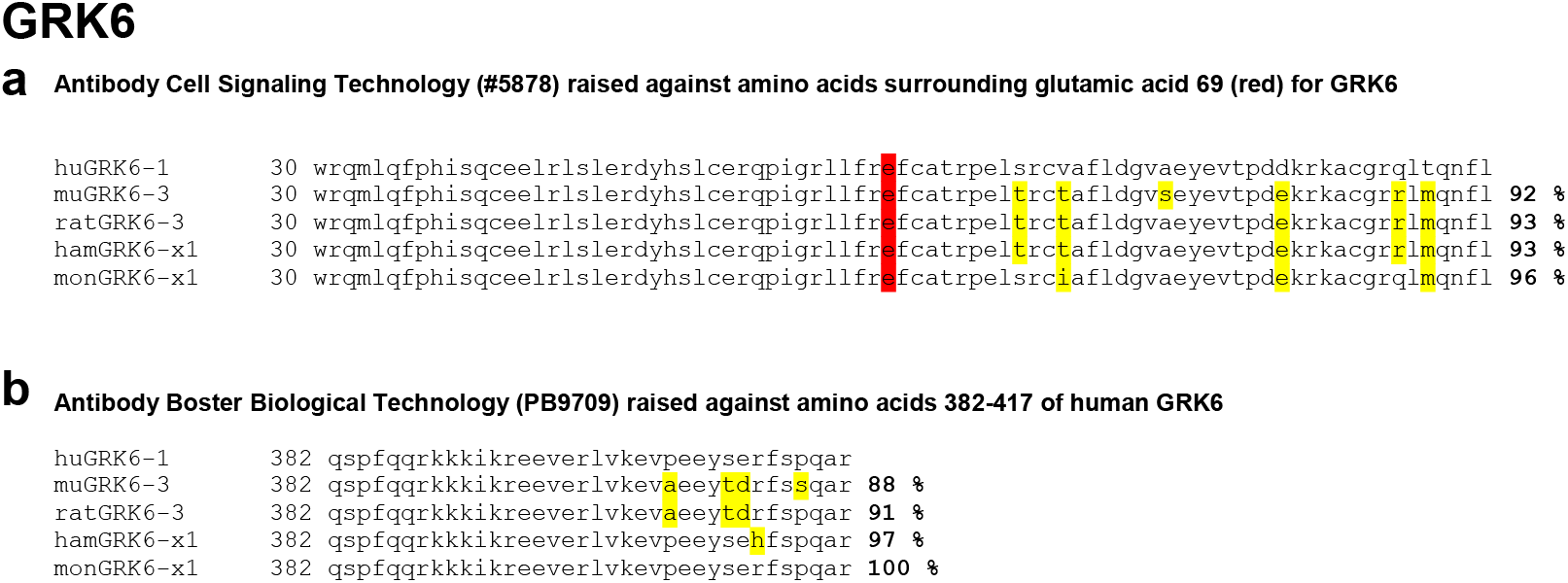
Alignment of human, mouse, rat, hamster, and monkey GRK6 protein sequences from amino acid 30 to 110 (**a**) and 382 to 417 (**b**). Differing amino acids are highlighted in yellow. Total identity is denoted in per cent.

**Supplementary Table 1.**
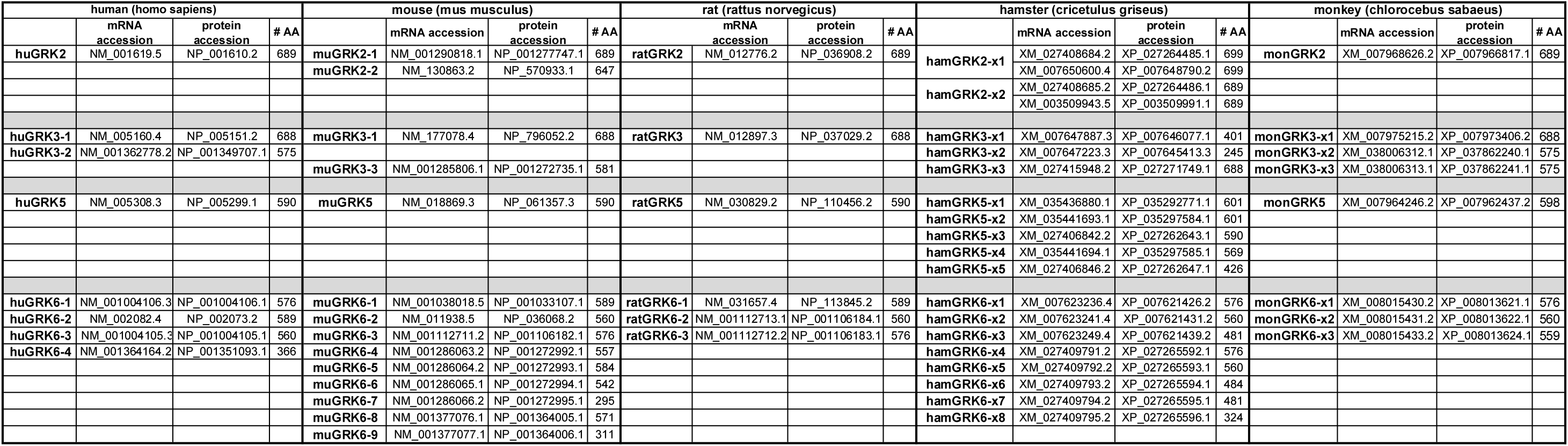
Last edited on 19th of October 2021. Overview of mRNA and protein accession identifiers of GRK isoforms in different mammals retrieved from https://www.ncbi.nlm.nih.gov. In case of human, mouse, and rat, the computationally predicted sequences reported are not shown, in case of hamster and monkey, all sequences shown are computationally predicted. Of note, some proteins within one species and GRK isoform are identical despite different accession numbers.

